# Coverage-versus-Length plots, a simple quality control step for *de novo* yeast genome sequence assemblies

**DOI:** 10.1101/421347

**Authors:** Alexander P. Douglass, Caoimhe E. O’Brien, Benjamin Offei, Aisling Y. Coughlan, Raúl A. Ortiz-Merino, Geraldine Butler, Kevin P. Byrne, Kenneth H. Wolfe

## Abstract

Illumina sequencing has revolutionized yeast genomics, with prices for commercial draft genome sequencing now below $200. The popular SPAdes assembler makes it simple to generate a *de novo* genome assembly for any yeast species. However, whereas making genome assemblies has become routine, understanding what they contain is still challenging. Here, we show how graphing the information that SPAdes provides about the length and coverage of each scaffold can be used to investigate the nature of an assembly, and to diagnose possible problems. Scaffolds derived from mitochondrial DNA, ribosomal DNA, and yeast plasmids can be identified by their high coverage. Contaminating data, such as cross-contamination from other samples in a multiplex sequencing run, can be identified by its low coverage. Scaffolds derived from the bacteriophage PhiX174 and Lambda DNAs that are frequently used as molecular standards in Illumina protocols can also be detected. Assemblies of yeast genomes with high heterozygosity, such as interspecies hybrids, often contain two types of scaffold: regions of the genome where the two alleles assembled into two separate scaffolds and each has a coverage level *C*, and regions where the two alleles co-assembled (collapsed) into a single scaffold that has a coverage level 2*C*. Visualizing the data with Coverage-versus-Length (CVL) plots, which can be done using Microsoft Excel or Google Sheets, provides a simple method to understand the structure of a genome assembly and detect aberrant scaffolds or contigs. We provide a Python script that allows assemblies to be filtered to remove contaminants identified in CVL plots.

**100-word article summary:** We describe a simple new method, Coverage-versus-Length plots, for examining *de novo* genome sequence assemblies. These plots enable researchers to detect scaffolds that have unusually high or unusually low coverage, which allows contaminants, and scaffolds that come from atypical parts of the organism’s DNA complement, to be detected. We show that contaminants are common in yeast genomes sequenced in multiplex Illumina runs. We provide instructions for making plots using Microsoft Excel or Google Sheets, and software for filtering assemblies to remove contaminants. Contaminants can be detected and removed, even without knowing their source.

## Introduction

It is now easy and cheap to sequence a yeast genome using the Illumina platform. Genome sequencing projects are usually described as either resequencing projects or *de novo* assembly projects. In resequencing projects, Illumina reads from the strain to be studied are mapped onto a reference genome from the same species (e.g. *Saccharomyces cerevisiae* S288C), in order to find nucleotide polymorphisms or mutations. In *de novo* assembly projects, the Illumina reads from the strain to be studied are assembled without using a reference, to produce a set of contigs or scaffolds for the strain. *De novo* assembly is used in several specific situations: (i), where no suitable reference genome sequence exists (e.g., if the species has not been sequenced before); (ii), where the researcher chooses to ignore the reference (e.g., to explore the pan-genome in natural isolates of *S. cerevisiae*); (iii), where the strain being studied comes from an unknown species; or (iv), for mixed samples, such as metagenomics projects. For any newly-discovered yeast species, in order to publish a formal taxonomic description in the *International Journal of Systematic and Evolutionary Microbiology*, authors are now requested to include a (*de novo*) sequence assembly of its genome^1^.

One of the most widely-used assembly programs for *de novo* yeast genome sequences is SPAdes (Bankevich et al. 2012). SPAdes assembles genomes into contigs or scaffolds, depending on the type of Illumina data that is input (single reads, paired-end reads, or mate-pair reads). Contigs are contiguous stretches of assembled genome sequence. Scaffolds are groups of contigs, whose relative order and orientation is known, but which are separated by short gaps of unknown sequence (represented by poly-N regions in the scaffold sequence). A typical yeast genome sequencing project will generate about 1.5 gigabases of raw data (e.g., 10 million Illumina reads of 150 bp each), which can be assembled by SPAdes in a few hours on a standard laptop, giving more than 50x coverage of the genome.

As the capacity of Illumina machines has grown, it has become increasingly common that samples are sequenced in multiplex, i.e. genomic libraries are constructed from pooled samples, using a different index (or pair of indexes) for each sample, and then sequenced together in the same flowcell. For example, an Illumina HiSeq 4000 machine can sequence 96 yeast samples in multiplex in a single flowcell. It should be noted that the Illumina multiplex methodology uses up to four separate sequencing reactions to obtain data from a single flowcell spot containing a genomic DNA fragment: the library index(es) are sequenced in 1-2 reactions using different primers than are used to sequence one or both ends of the genomic DNA fragment^2^. There is therefore some potential for mix-ups in which genomic DNA reads are assigned to the wrong index and hence to the wrong sample.

The drawback to our ability to sequence genomes faster and cheaper is that the amount of time and care spent on human examination of each sequence must inevitably decrease (Lu and Salzberg 2018). Gross errors such as the assignment of a genome sequence to the wrong species (Stavrou et al. 2018; Watanabe et al. 2018; Shen et al. 2016; Pavlov et al. 2018), or yeast genome sequences that include many contigs from contaminating bacteria (Donovan et al. 2018), are increasingly present in public databases. In addition, the genome sequences of some yeast isolates have proven difficult to assemble because they are highly heterozygous, in some cases because they are interspecies hybrids (Pryszcz et al. 2015; Pryszcz and Gabaldon 2016; Schröder et al. 2016; Braun-Galleani et al. 2018). In this report, we present a simple method that can be used for quality control of an Illumina genome assembly, by making use of the information that SPAdes produces about the length and coverage of each contig or scaffold.

## Materials and Methods

All the example assemblies we present were made using SPAdes version 3.11.1 (Bankevich et al. 2012), downloaded from http://cab.spbu.ru/software/spades/. Illumina FASTQ files were obtained by our laboratory using commercial DNA sequencing services, except for *Hanseniaspora osmophila* NCYC58 for which we downloaded FASTQ files from http://opendata.ifr.ac.uk/NCYC/ and assembled them (Table 1). We did not carry out any data preprocessing steps.

**Table 1.** Summary of Illumina datasets analyzed, and their SPAdes assemblies.

| Species | Strain | Type of Illumina data<sup>a</sup> | Million reads | Scaffold N50<sup>b</sup> | Scaffold L50<sup>b</sup> | Number of scaffolds | Scaffolds > 500 bp | Modal *k*-mer coverage | Minimum scaffold size |
| --- | --- | --- | --- | --- | --- | --- | --- | --- | --- |
| *Komagataella phaffii* | CBS7435 | PE, 150 bp | 11.1 | 642 kb | 5 | 24760 | 614 | 73x | 78 bp |
| *Hanseniaspora osmophila* | NCYC58 | PE, 100 bp | 13.8 | 71 kb | 51 | 2692 | 474 | 35x | 56 bp |
| *Torulaspora delbrueckii* | L17 | SE, 50 bp | 20.8 | 4 kb | 392 | 27260 | 4507 | 33x | 34 bp |
| *Torulaspora delbrueckii* | L16 | SE, 50 bp | 18.9 | 23 kb | 123 | 12869 | 6721 | 29x | 34 bp |
| *Metschnikowia* aff. *pulcherrima* | UCD127 | SE, 150 bp | 6.7 | 2 kb | 3226 | 39964 | 14981 | 29x | 78 bp |

Assembly statistics were calculated using QUAST from scaffolds >500 bp.
<sup>a</sup> PE, paired end reads. SE, single reads.
<sup>b</sup> When scaffolds are sorted in descending order of length, N50 is the length of the scaffold that contains the midpoint base in the assembly, and L50 is the number of this scaffold.

Instructions for how to make a CVL plot from a SPAdes output file using common spreadsheet programs are given in Box 1.

## Results

SPAdes (Bankevich et al. 2012) is a popular assembly program because it is simple to install, runs in a few hours on a laptop, and makes good assemblies ‘out of the box’ with its default settings. It produces two main output files, called `contigs.fasta` and `scaffolds.fasta`. The `scaffolds.fasta` file may include some poly-N regions where two contigs have been joined with a gap between them, because their relative order and orientation is known. For single-read and paired-end reads data, the two files are very similar and we consider only the `scaffolds.fasta` output file here.

The `scaffolds.fasta` file contains all the assembled scaffolds that are larger than a minimum size. The minimum size is *k*+1 bp, where *k* is the largest *k*-mer size that SPAdes used for the assembly. For 150-bp reads, *k* is usually 77 bp so the smallest scaffolds reported are only 78 bp long. In the file, the scaffolds are given names such as NODE_1, NODE_2, etc, and are sorted in decreasing order of length so that NODE_1 is the longest scaffold in the assembly. Each scaffold in the file has a FASTA header that includes its name, its length in basepairs, and its coverage (Fig. 1).

**Figure 1.**
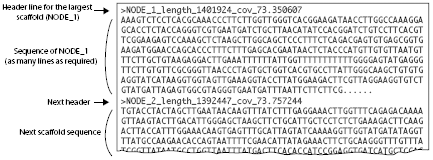
Organization of a `scaffolds.fasta` file containing a SPAdes genome assembly. The header line for each scaffold begins with a “>” character, which is standard for data in FASTA format. The header for each scaffold shows the scaffold’s node number, its length, and its *k-mer* coverage. In this example, NODE_1 is 1,401,924 bp long and has 73.35x coverage. The header line is followed by NODE_1’s DNA sequence, and then the header and sequence of the next scaffold (NODE_2). The file can contain thousands of scaffold sequences, one after the other, down to the minimum scaffold size. The scaffolds are always named and sorted in descending order of length.

The coverage values reported by SPAdes are properly called *k*-mer coverage. *k*-mer coverage is directly proportional to the more familiar concept of base coverage, which is the average number of Illumina reads that covered each nucleotide site in the scaffold when its consensus sequence was generated. The two quantities are related by the formula^3^

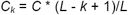

where *C*_*k*_ is *k*-mer coverage, *C* is base coverage, *L* is the read length, and *k* is the largest *k*-mer size that was used by SPAdes to make the assembly. *C*_*k*_ for a scaffold is the average number of times the *k*-mers that make up the scaffold were seen among the reads. For 150-bp reads and *k*-mers of 77 bp, the *k*-mer coverage is approximately half the base coverage.

### Example 1: A typical yeast Coverage-versus-Length (CVL) plot

In this example, we used a commercial genome sequencing service to sequence a yeast strain constructed from *Komagataella phaffii* strain CBS7435 (this species was formerly called *Pichia pastoris*). Two high-quality reference genome sequences are available for CBS7435 (Sturmberger et al. 2016; Love et al. 2016). Our raw data consisted of 11.1 million paired-end reads of 150 bp each. SPAdes assembled the data into almost 25,000 scaffolds, but most of these are very short and only 614 scaffolds are larger than 500 bp (Table 1). The N50 of the assembly is 642 kb. Researchers would typically discard very short scaffolds from the assembly, but here we retained them for analysis.

Figure 2A shows a coverage-versus-length (CVL) plot for this assembly. To make a CVL plot, we simply extract the coverage and length data from all the headers in the `scaffolds.fasta` file (Fig. 1), and plot them against each other. Each point in the plot is a scaffold. The X-axis is scaffold length, plotted on a linear scale. The Y-axis is the coverage (*k*-mer coverage), plotted on a log scale. The longest scaffold in the *K. phaffii* assembly (NODE_1) is 1.4 Mb long and has a coverage of 73x. The other large scaffolds (>100 kb) have similar levels of coverage (between 70x and 90x). The variation in coverage levels increases on shorter scaffolds. As expected, all the large scaffolds matched the reference nuclear genome sequence available for *K. phaffii*. A few scaffolds stand out because they have coverage that is much higher than the 70-90x seen in the large scaffolds from the nuclear genome. NODE_19 is the mitochondrial genome (36 kb, 1493x). NODE_26 is a cytoplasmic linear DNA plasmid related to killer plasmids (12 kb, 557x). Coverage levels vary widely in the smaller scaffolds, shown in a zoomed-in view in Figure 2B. Five small scaffolds with very high coverage (1600x – 3000x) come from the ribosomal DNA array. Thus, scaffolds that stand out on the basis of unusually high coverage are those that do not come from typical parts of the nuclear genome.

**Figure 2.**
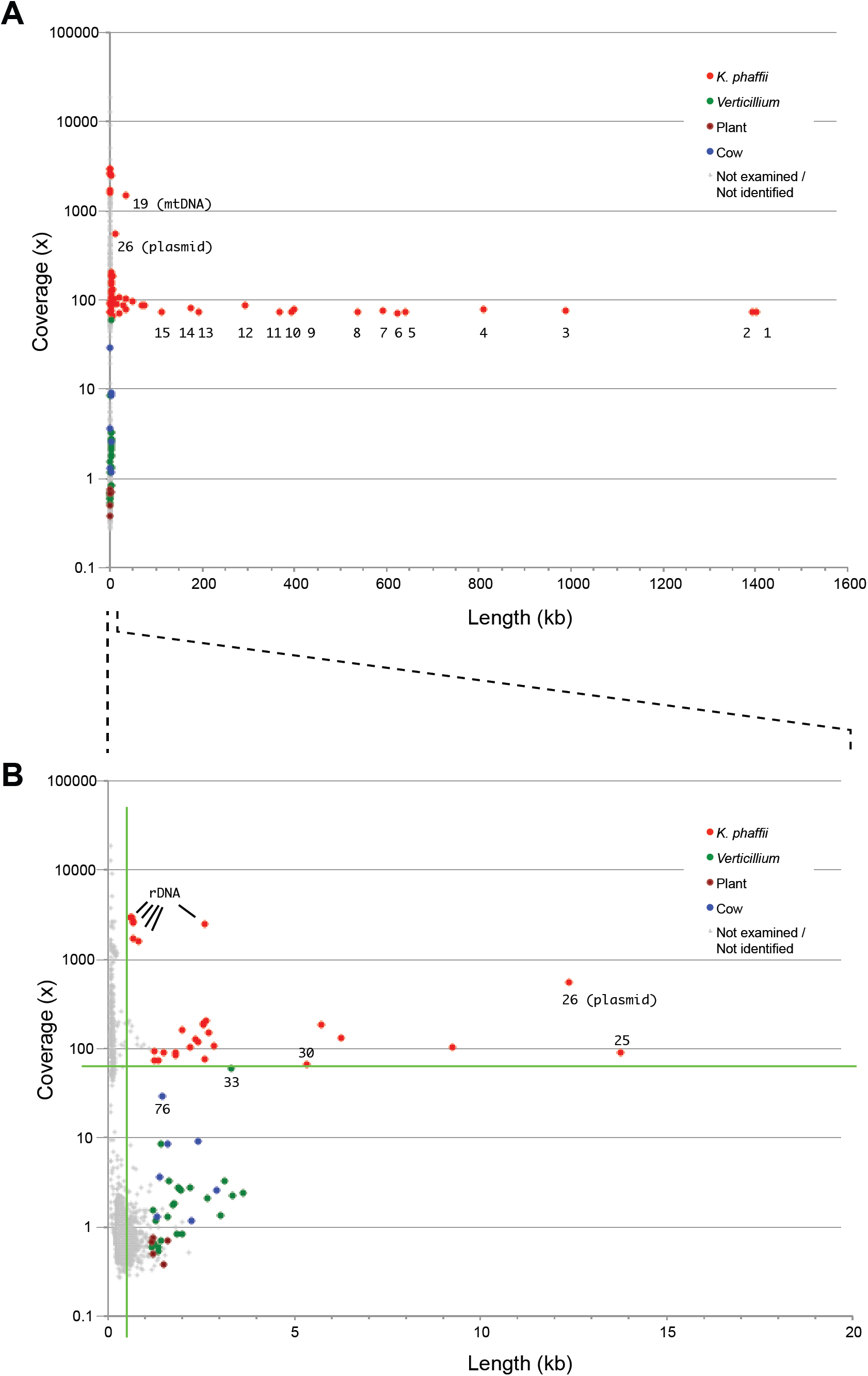
CVL plot of a SPAdes assembly of data from *Komagataella phaffii* strain CBS7435, showing (A) all scaffolds, and (B) scaffolds shorter than 20 kb. The sources of the longest 100 scaffolds were identified using BLASTN searches against the NCBI database and are color coded. Numbers beside points indicate scaffold (NODE) numbers mentioned in the text. The green lines in B indicate 65x coverage and 500 bp length.

We used BLASTN searches against the NCBI nonredundant nucleotide database to identify the origins of each of the 100 longest scaffolds (i.e., all the scaffolds longer than 1160 bp), and found that none of the ones with low coverage come from *K. phaffii* (Fig. 2B). The most common source of contamination (25 scaffolds) was *Verticillium dahliae* (a fungal pathogen of plants), including its nuclear rDNA sequence (NODE_33, 3326 bp at 60x coverage) as well as multiple scaffolds containing parts of its mitochondrial genome (∼3x coverage) and a nuclear retrotransposon (∼1x). We also identified seven contaminating scaffolds from cow (all were nuclear satellite DNAs, of which the most abundant was NODE_76, corresponding to the bovine ‘1.715’ satellite, 1397 bp at 29x coverage), and five from plant species (nuclear repetitive DNA and chloroplast DNA). We suspect that the source of these diverse contaminating sequences was other sequencing projects being carried out at the commercial sequencing center, probably in multiplex on the same Illumina flowcell as our sample. We do not think that our *K. phaffii* assembly is unusually highly contaminated, but rather that the situation we describe is commonplace.

A bioinformatician handling our *K. phaffii* assembly would normally discard the shorter scaffolds before analysis or submission of the data to public databases. Typical practice would be to discard scaffolds < 500 bp or < 1 kb. However, as Figure 2B shows, our assembly includes some scaffolds up to 4 kb long that are contaminants. The CVL plot allows the existence of scaffolds with aberrant coverage levels to be detected. In this example, applying a filter to keep only scaffolds with coverage >63x and length >500 bp would exclude all the contaminants and retain almost all the genuine *K. phaffii* scaffolds (green lines in Fig. 2B). Importantly, these cutoffs could have been chosen simply by looking at the CVL plot and using BLASTN to check the most anomalous scaffolds (NODES 76, 33, and 30; Fig. 2B).

### Example 2: Aneuploidy, and bacteriophage contamination

Figure 3 shows a CVL plot of a SPAdes assembly from *Hanseniaspora osmophila* strain NCYC58. This strain was sequenced by the UK National Collection of Yeast Cultures at the Earlham Institute as part of a systematic strain sequencing program (Dujon and Louis 2017; Wu et al. 2017), and assembled by us (Table 1). A high-quality whole-genome shotgun assembly of a reference *H. osmophila* strain (AWRI3579) is also available (Sternes et al. 2016).

**Figure 3.**
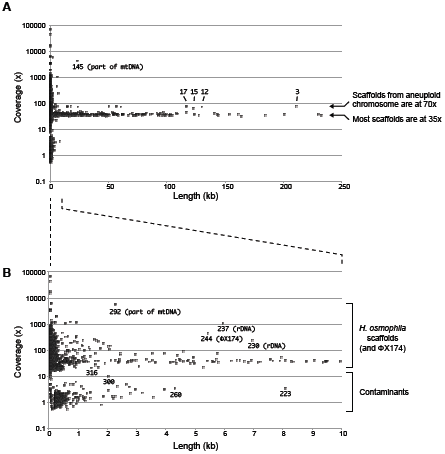
CVL plot of a SPAdes assembly of data from *Hanseniaspora osmophila* strain NCYC58 showing (A) all scaffolds, and (B) scaffolds shorter than 10 kb. Numbers beside points indicate scaffold (NODE) numbers mentioned in the text.

The plot of the NCYC58 assembly shows many large scaffolds with coverage of ∼35x (Fig. 3A). It also shows many small scaffolds with coverage <10x (Fig. 3B), which in this case are contamination from other yeast species (probably due to cross-contamination from multiplexed yeast samples). From BLASTN searches, we found that the largest of these contaminating scaffolds (NODE_223, 8087 bp, 3x coverage) comes from *Saccharomyces cerevisiae* mitochondrial DNA. The second-largest (NODE_260, 4308 bp, 3x coverage) comes from *Wickerhamomyces anomalus* mitochondrial DNA. The most abundant yeast contaminant (NODE_300, 2033 bp, 9x coverage) comes from *S. cerevisiae* ribosomal DNA. There is a clear gap in coverage between the contaminants and the least-abundant genuine scaffold from *H. osmophila* (NODE_316, 1472 bp, 21x coverage), so the contaminating yeast scaffolds could be removed by excluding scaffolds with <15x coverage (Fig. 3B).

The CVL plot for NCYC58 also shows evidence of aneuploidy in this strain. Although most of the large scaffolds have coverage ∼35x, some scaffolds have twofold higher coverage at ∼70x (for example, NODES 3, 12, 15 and 17 in Fig. 3A). By BLASTN searches against the reference sequence of *H. osmophila* (Sternes et al. 2016), we found that the NCYC58 scaffolds with ∼70x coverage all map to the same chromosome in the reference assembly (AWRI3579 scaffold10). Therefore, we suggest that there is an extra copy of this chromosome in strain NCYC58.

The high-coverage scaffolds in the CVL plot for NCYC58 include the mitochondrial genome and the ribosomal DNA, as was seen in *K. phaffii*. Another high-coverage scaffold is a contaminant: the bacteriophage ΦX174 genome (NODE_244, scaffold length 5441 bp, coverage 477x). ΦX174 DNA is used as an internal positive control for the sequencing reagents in some Illumina protocols and is a frequent contaminant in microbial genome assemblies (Mukherjee et al. 2015; Lu and Salzberg 2018). In CVL plots from some other yeast genome assemblies (not illustrated here) we have also found the bacteriophage λ genome at high coverage (scaffold length 48-49 kb). Bacteriophage λ DNA is used as a standard for DNA quantification in an Illumina protocol for making indexed paired-end libraries^4^.

### Example 3: Assemblies of highly heterozygous or hybrid genomes

Figure 4A shows the CVL plot of a SPAdes genome assembly from *Torulaspora delbrueckii* strain L17. The plot shows that two major types of scaffold are present in L17: ones with coverage ∼33x, and ones with coverage ∼16x. Both types of scaffold have high similarity to other *Torulaspora* sequences in databases, so they are not contaminants. By BLASTN searches against the genome sequence of the type strain of *T. delbrueckii* (CBS1146; Gordon et al. 2011) we found that L17 is a highly heterozygous diploid strain. Heterozygosity is unusual, because most strains of *T. delbrueckii* are haploid and their assemblies do not show two types of scaffold (e.g. strain L16 in Fig. 4B).

**Figure 4.**
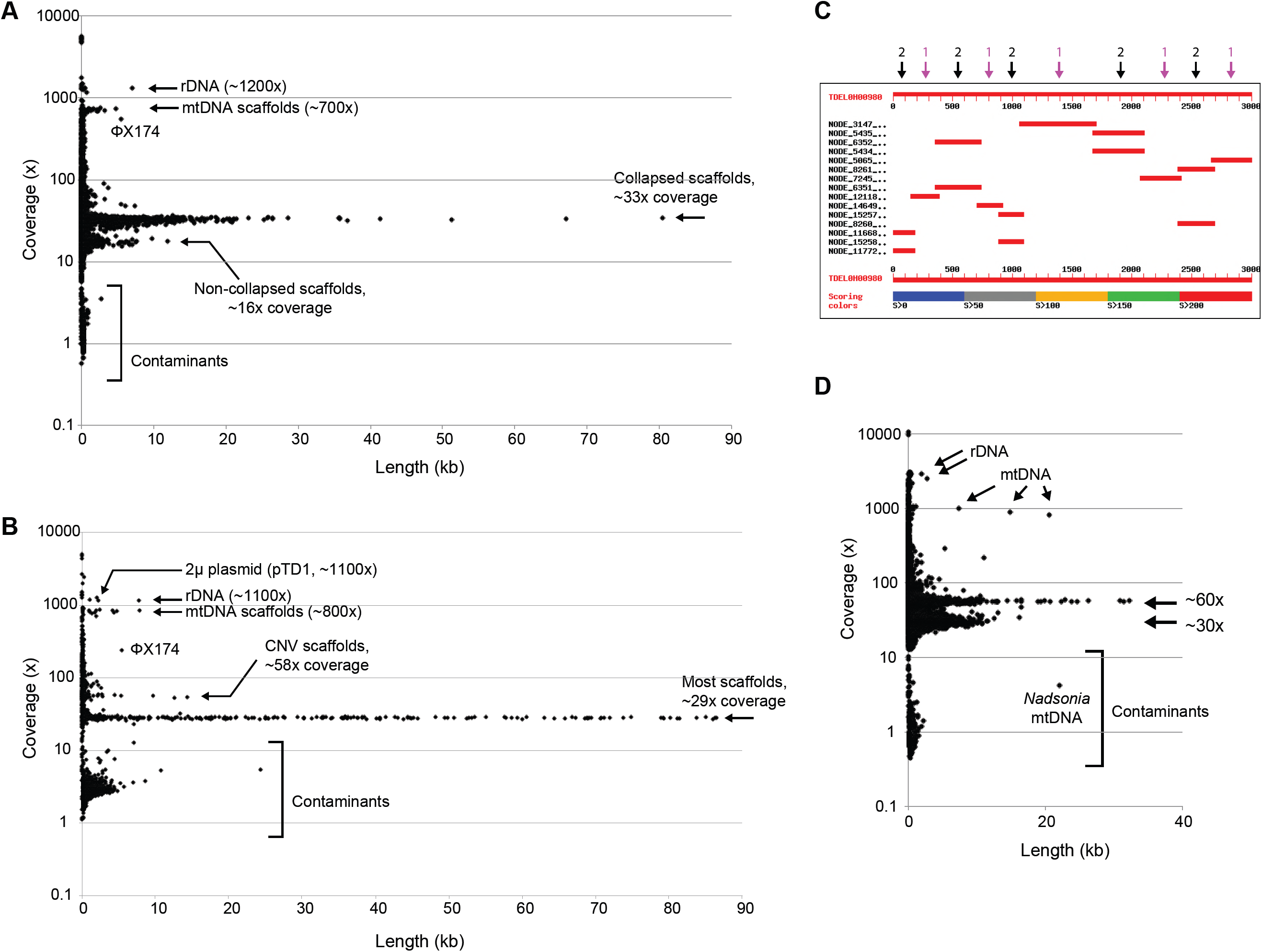
Highly heterozygous genomes. (A) CVL plot of the assembly from *Torulaspora delbrueckii* strain L17, which is highly heterozygous. (B) CVL plot of the assembly from *T. delbrueckii* strain L16, which is not heterozygous. (C) Graphical view of a BLASTN search result. The database is the assembly of the L17 genome, and the query is the first 3 kb of the *DYN1* gene of the *T. delbrueckii* reference strain CBS1146. Each horizontal red bar indicates a scaffold that contains a region of *DYN1*, relative to the 3-kb scale. Black arrows mark allelic pairs of scaffolds from non-collapsed regions, and magenta arrows mark single collapsed scaffolds. For clarity, this diagram shows only the first 3 kb of the 12 kb *DYN1* gene. In total, *DYN1* maps onto 68 scaffolds in the L17 assembly: 22 collapsed scaffolds and 23 pairs. (D) CVL plot of the assembly from *Metschnikowia* strain UCD127.

Detailed analysis shows how the SPAdes assembler produced this pattern from the L17 data. In some sections of the genome, SPAdes ‘collapsed’ (merged) the sequences of the two alleles into a single consensus sequence, making only one scaffold, with ∼33x coverage. In other sections of the genome, SPAdes kept the two alleles separate, resulting in two scaffolds from these regions, with ∼16x coverage each. The two types of sections alternate along the genome, and some of the sections are quite small – only a few hundred basepairs. For example, a BLASTN search using part of the large gene *DYN1* as the query, and the SPAdes assembly of the L17 genome as the database, produces hits with the pattern shown in Fig. 4C, alternating between hits to single scaffolds, and hits to pairs of scaffolds with lower coverage. Strain L17 contains two copies (alleles) of *DYN1*, but the assembler generated 68 separate scaffolds from them. In the parts of *DYN1* where separate allelic scaffolds were produced, the sequence divergence between alleles is approximately 7%. One of the alleles is almost identical to the *T. delbrueckii* CBS1146 reference genome, while the other is almost identical to the allele present in strain L16. Similar results are seen when other genes or genomic regions are used as BLASTN queries against the L17 assembly.

The CVL plot for UCD127, a strain from the *Metschnikowia pulcherrima* subclade that we isolated from nature, shows a similar pattern with two major types of scaffold with a twofold difference in their coverage levels (Fig. 4D). UCD127 is either a highly heterozygous diploid or an interspecies hybrid (Venkatesh et al. 2018). In this example, a large contaminant scaffold (22 kb, 4x coverage) was the mitochondrial genome of *Nadsonia starkeyi-henricii*, a yeast that we sequenced in the same multiplex job (O’Boyle et al. 2018), which demonstrates that cross-contamination occurred in the flowcell.

## Discussion

CVL plots provide a simple way to detect scaffolds or contigs in an assembly that have aberrant levels of sequence coverage. In some cases these scaffolds represent DNA that is truly present in the organism targeted for sequencing, at a copy-number that is higher or lower than average, which allows CVL plots to be used to detect mitochondrial DNA, ribosomal DNA, plasmids or other extrachromosomal elements, and aneuploidy. In other cases, the scaffolds with unusual coverage represent contaminants, which can stem from several different sources: cross-contamination from other DNA samples sequenced in the same multiplex experiment; bacterial contamination in the DNA sample used for sequencing; and bacteriophage DNA deliberately introduced for quality control in the sequencing process.

In most of the examples we presented, we used assemblies from yeast strains where a reference genome assembly was already available from the same species. We did this so that we could validate our interpretations of the CVL plots by using BLAST searches. However, it should be emphasized that CVL plots allow most contaminant scaffolds to be detected and removed, even when no reference is available for the genome being sequenced, and even when the contaminant’s source species is not in genome databases. Contaminants can be detected and removed even without knowing what they are.

The low-coverage contaminants we detected correspond to sequences that are probably the highest copy-number sequences in their source genomes: ribosomal DNAs and mitochondrial DNAs (e.g., from *Verticillium*), satellite DNAs (bovine), retrotransposons (*Verticillium*), and chloroplast DNA (plants). This result is to be expected if cross-contamination occurs in a multiplex sequencing run; even a small number of sequencing reads from a eukaryotic genome can be assembled into consensus sequences for the most abundant repeat families in that genome (Neuveglise et al. 2002). As sequencing costs fall, the coverage level to which genomes are sequenced tends to increase. But as coverage increases, the lengths of scaffolds assembled from contaminating DNA will also increase.

We have illustrated CVL plots with examples that used the SPAdes assembler, but in principle CVL plots could be made using the output from any assembler that reports coverage levels for individual scaffolds or contigs. SPAdes is particularly well suited to CVL plots because one of its characteristic behaviors is that contigs terminate whenever they enter repeat sequences, so that repeat sequences and the unique sequences that flank them are reported as separate contigs. This behavior means that each contig or scaffold reported by SPAdes has a fairly uniform level of coverage along its length.

A researcher who has assembled a genome sequence will usually want to ‘clean up’ the assembly before submitting it to a public database. The current standard cleanup step is to remove all scaffolds smaller than a cutoff length, which is usually 500 bp or 1 kb. However, as we have shown, many contaminants would still be retained after such a length filter alone. Even without doing any BLAST searches, a CVL plot will enable the researcher to choose a cutoff value for minimum coverage. Applying cutoffs to both coverage and length will allow the researcher to produce an assembly whose level of contamination is greatly reduced. CVL plots can be made from SPAdes assemblies by using spreadsheet software without any programming steps (Box 1). We also provide a simple Python script^5^ that researchers can use to filter a SPAdes `scaffolds.fasta` or `contigs.fasta` file so that only the scaffolds/contigs that have coverage and length above minimum values specified by the user are retained.

## Acknowledgements

This work was supported by the Wellcome Trust PhD programme in Computational Infection Biology at University College Dublin (105341/Z/14/Z). We thank Caroline Wilde (Lallemand, Inc.) for strains L16 and L17. The NCYC58 sequence data were produced by the National Collection of Yeast Cultures (UK), in partnership with the Earlham Institute, using funding awarded to the Institute of Food Research, Norwich, UK by the Biotechnology and Biological Sciences Research Council.

### Box 1. How to make a CVL plot using Microsoft Excel, Google Sheets, or R.

**(A) Instructions for Excel.** These instructions are based on Excel for Mac 2011. Other versions of Excel may be slightly different. A video demonstration of these instructions is available at https://tinyurl.com/CVLexcel

1. 1.Import the `scaffolds.fasta` file into a new blank Excel sheet:
  - In Excel, choose File → Open → `scaffolds.fasta` (Click Next and Finish in the dialog box).
  - Select Column A.
  - Choose Data → Sort → Sort by Column A (in A-to-Z order). All the header lines go to the top of the Excel sheet, with all the sequence rows below them.
  - Delete all the sequence data: Find the first row of sequence data and click on its row. Then scroll to the bottom of the file and shift-click on the last row of sequence data. Then choose Edit → Delete.
  - You are left with all the header lines. Save the file as an Excel (.`xlsx`) file.
2. 2.Split the information in the header lines into separate columns:
  - Choose Data → Text to Columns. Click Next once.
  - In the Delimiters box, check Other and type an underscore character (_) in the box. Then click Next and Finish.
  - You should see the scaffold length in column D, and the coverage data in column F.
3. 3.Make a scatterplot:
  - Select column D and command-click to select column F as well.
  - Insert → Chart… → Scatter → Marked Scatter
  - Change the Y-axis (coverage) to a log scale by double-clicking on the Y-axis and ticking the “Logarithmic scale” checkbox.
  - You can identify the scaffold associated with any point by moving the mouse over it.
  - It’s easier to see individual points if you make them smaller: Double-click on any point, choose Marker Style, and reduce the marker size to 3 point.

**(B) Instructions for Google Sheets.** A video demonstration of these instructions is available at https://tinyurl.com/CVLgsheets

1. Rename the `scaffolds.fasta` file `scaffolds.txt`
2. Import the `scaffolds.txt` file into a new blank Google Sheet:
  - Choose File → Import… → Upload, and select the file `scaffolds.txt`
  - Click Import Data (leave all the options as their default values), and wait while the data is imported (it can take 1-2 minutes).
  - Select Column A.
  - Choose Data → Sort Sheet by Column A, A→Z, and wait while the data is sorted. All the header lines go to the top of the sheet, with all the sequence rows below them. Wait until the progress bar on the top right has finished.
  - Delete all the sequence data: Find the first row of sequence data and click on its row. Then scroll to the bottom of the file and shift-click on the last row of sequence data. Then choose Edit → Delete Rows.
  - You are left with all the header lines.
3. Split the information in the header lines into separate columns:
  - Select Column A.
  - Choose Data → Split Text into Columns…
  - A box labeled Separator pops up near the bottom of the screen. Pull down the menu in this box, choose Custom, and type an underscore character (_) in the box. Then hit Enter on the keyboard.
  - You should see the scaffold length in column D, and the coverage data in column F.
4. Make a scatterplot:
  - Select column D and control-click to select column F as well.
  - Insert → Chart…
  - In the Chart Editor panel (on the right of the screen), under the Data tab, choose Chart Type → Scatter
  - In the Chart Editor panel, under the Customize tab, click on Vertical Axis and tick the checkbox beside Logarithmic Scale.
  - You can identify the scaffold associated with any point by clicking on the point.
  - It’s easier to see individual points if you make them smaller. In the Chart Editor, under the Customize tab, choose Series, and reduce the Point size to 2 px.

**(C) Instructions for R.** We provide an R script for making CVL plots at https://github.com/APDLS/CVLFilter

## Footnotes

1 http://ijs.microbiologyresearch.org/content/journal/ijsem/about

2 https://support.illumina.com/downloads/indexed-sequencing-overview-15057455.html

3 https://www.ebi.ac.uk/~zerbino/velvet/Manual.pdf

4 https://support.illumina.com/content/dam/illumina-support/documents/documentation/chemistry_documentation/samplepreps_trusight/trusight-rapid-capture-reference-guide-15043291-01.pdf

5 https://github.com/APDLS/CVLFilter

